# Effectiveness of Growth Hormone–Releasing Hormone Agonists (GHRH-A) in Chronic Kidney Disease-Induced Heart Failure with Preserved Ejection Fraction

**DOI:** 10.1101/2020.05.30.111476

**Authors:** Angela C. Rieger, Luiza L Bagno, Alessandro Salerno, Victoria Florea, Jose Rodriguez, Marcos Rosado, Darren Turner, Lauro M. Takeuchi, Raul Dulce, Wayne Balkan, Ivonne H. Schulman, Andrew Schally, Joshua M. Hare

## Abstract

**Background:** Therapies that improve morbidity and mortality in heart failure with preserved ejection fraction (HFpEF) are lacking. Growth hormone releasing hormone analogues (GHRH-A) reverse fibrosis and improve cardiac function in ischemic and non-ischemic animal models. We tested the hypothesis that GHRH-A treatment ameliorates chronic kidney disease (CKD)-induced HFpEF in a large animal model.

**Methods:** Female Yorkshire pigs (n=16) underwent 5/6 nephrectomy via renal artery embolization, which induced HFpEF, and 12-weeks later received daily subcutaneous injections of GHRH-A (n=8) or placebo (n=8). Kidney function, renal and cardiac MRI, pressure-volume loops, and electrical stimulation were assessed at baseline, 12-weeks, and 16-18 weeks post-embolization.

**Results:** The CKD model was confirmed by increased creatinine and BUN. HFpEF was demonstrated at 12 weeks by maintenance of ejection fraction associated with increased left ventricular mass, relative wall thickening, end-diastolic pressure (EDP), end-diastolic pressure-volume relationship (EDPVR), and tau. After 6 weeks of treatment, diastolic function improved in the GHRH-A group, evidenced by normalization of EDP (p=0.03) associated with improved diastolic compliance as measured by EDP/EDV ratio (p=0.018).

**Conclusion:** A beneficial effect of GHRH-A in diastolic function was observed in a CKD large animal model that manifests the characteristics of HFpEF. These findings have important therapeutic implications for the HFpEF syndrome.

## INTRODUCTION

The prognosis for heart failure with preserved ejection fraction (HFpEF), a complex disease involving multiple comorbidities with limited treatment options^1–3^, has not improved over decades. Particularly, chronic kidney disease (CKD)-associated HFpEF has worse outcomes than other HFpEF phenotypes^4^. It manifests predominantly with pathologic left ventricular (LV) remodeling, characterized by LV hypertrophy with increased LV-mass index and relative wall thickness (RWT), impaired ventricular-arterial coupling and right ventricular relaxation, and increased stroke work (SW)^4^.

Growth hormone-releasing hormone (GHRH) is a pleiotropic hormone that has extra-pituitary effects. In murine models of ischemic cardiomyopathy (ICM) with reduced ejection fraction^5^, GHRH-agonists (GHRH-A) improved cardiac function and decreased scar size, mediated by myocardial GHRH-receptors^6 7^ Further mechanisms of action included cardiac progenitor proliferation and decreased cardiomyocyte apoptosis and inflammatory responses^6^. GHRH-A (MR-409) also improved diastolic strain and reduced myocardial scar in a swine large animal model with subacute ICM^8^. Notably, in a HFpEF murine model using chronic administration of angiotensin, GHRH-A prevented impairment of cardiomyocyte contractile function and relaxation and development of HFpEF features, including LV hypertrophy, collagen deposition, and diastolic dysfunction, compared to the placebo^9^. In this study, we used a swine large animal model of CKD-induced HFpEF that displays the characteristics observed in humans to test the hypothesis that GHRH-A ameliorates the cardiac diastolic dysfunction.

## METHODS

The study was conducted in a blinded fashion. All animal protocols were approved by the University of Miami Institutional Animal Use and Care Committee.

Female Yorkshire swine (30-35kg) underwent catheter-induced embolization with polyvinyl alcohol particles^10, 11^ (150 −250μ Boston Scientific) and 100% ethanol infusion to produce the 5/6 nephrectomy CKD model. Post-embolization arteriogram was performed to assess the residual renal flow to the embolized and remnant kidney. Twelve weeks post-embolization, animals were randomized to receive daily subcutaneous injections of GHRH-A (n=8, MR-409) or placebo (n=8, Plasma-Lyte, Baxter, IL, USA).

### Magnetic Resonance Imaging

Renal anatomical structure and perfusion was evaluated by renal magnetic resonance imaging (MRI, 3.0T clinical scanner, Magnetom, Siemens AG, Munich, Germany). Renal function was assessed by glomerular filtration rate (GFR) measurement using Iohexol and other laboratory measurements. Follow up evaluations were done weekly in the initial month and monthly throughout the GHRH-A treatment or placebo. Animals were studied for a total of 16-18 weeks in a blinded manner.

Cardiac MRI was conducted at the Interdisciplinary Stem Cell Institute to evaluate cardiac function (Siemens Trio 3T Tim, Erlangen, Germany) with Syngo MR Software using a 16-channel body surface coil with ECG gating and short breath-hold acquisitions. Cardiac evaluation included ECG-gated cine, first-pass gadolinium delayed hyperenhancement images, and tagging^12^.

### Hemodynamic Assessment

Pressure-Volume (PV) Loops were done at baseline, 12 weeks post-embolization, and before sacrifice with a PV catheter (VENTRI CATH 507for Large Animals)^13^. After Seldinger technique, the catheter was placed within the left ventricle and steady state PV loops were performed. Subsequently, the inferior vena cava was occluded with a balloon to determine end-diastolic pressure-volume relationship (EDPVR) and end-systolic pressure-volume relationship (ESPVR). Data were acquired after the animals were given time to stabilize.

### Electrophysiologic Assessment

Programmed Electrical Stimulation (PES) pacing was performed at twice diastolic threshold using an electro stimulator triggered by a personal computer, via a bipolar catheter placed intravenously in the right ventricular apex and left ventricular apex. For a refractory period assessment, a drive train sequence of 8 to10 pulses (S1) is delivered at S1S1 cycle length 400 ms, followed by the delivery of premature extra stimuli (S2) at 340 ms, 320 ms, and reduced in 20-ms decrements to the point where the S2 impulse does not capture, i.e., at the ventricular effective refractory period (VERP). The Induction of Sustained Ventricular Arrhythmias was performed in each animal at 12 week and euthanasia time points after embolization. This stimulation was performed in the right ventricle and left ventricle.

### Calcium Measurements

Cardiac myocytes were isolated and prepared from heart biopsies of swine by adapting a protocol described previously^14^. Briefly, left ventricular biopsies were collected from the swine at the operative room immediately before sacrifice. Specimens were kept in an ice-cold cardioplegic buffer (in *mmol/L*: 50 KH_2_PO_4_, 8 MgSO_4_, 10 HEPES, 5 adenosine, 140 glucose, 100 mannitol, 10 taurine) and processed within 10 minutes. Myocardial tissues were washed twice and then minced into small pieces (~3mm). Tissue chunks were transferred into a digestion device and then subjected to sequential (6 steps) enzymatic digestion (at 37°C) using collagenase type V (Worthington Biochemical Corporation, Lakewood, NJ) 250 U/mL and protease type XXIV (Sigma-Aldrich Co., Saint Louis, MO) 4 U/mL, in a digestion buffer containing (in *mmol/L*): 1.2 MgSO_4_, 10 glucose, 20 taurine, 113 NaCl, 4.7 KCl, 0.6 KH_2_PO_4_, 0.6 NaH_2_PO_4_, 12 NaHCO_3_, 12 KHCO_3_, 4 Na-pyruvate, 10 butadiene monoxime). Supernatants containing the cardiomyocytes were collected in digestion buffer with 0.5 % bovine serum albumin (BSA, Sigma-Aldrich Co. Saint Lois, MO). Then, the cells were subjected to a sequential extracellular Ca^2+^ restoration on a Tyrode’s buffer containing (in *mmol/L*): 144 NaCl, 1 MgCl_2_, 10 HEPES, 5.6 glucose, 5 KCl, 1.2 NaH_2_PO_4_ (adjusted to a pH 7.4 with NaOH). Finally, cardiomyocytes were resuspended in Tyrode’s buffer containing 1.8 mmol/L CaCl2 at room temperature until being used.

Intracellular Ca^2+^ was measured using the Ca^2+^-sensitive dye Fura-2 and a dual-excitation (340/380 nm) spectrofluorometer (IonOptix LLC, Milton, MA, USA). First, cardiomyocytes were incubated with 2.5 μmol/L Fura-2 for 15 minutes at room temperature and then washed with fresh regular Tyrode’s solution for at least 10 minutes. Then, cardiomyocytes were placed in a perfusion chamber adapted to the stage of an inverted Nikon eclipse TE2000-U fluorescence microscope. Cells were superfused with a Tyrode’s buffer at 37°C and Fura-2 fluorescence was acquired at an emission wavelength of 515±10 nm. The cardiomyocytes were electric field-paced (20V) at different frequencies from 0.2 to 1.5 Hz. The calibration was performed in cardiomyocytes “ex vivo” superfusing a free Ca^2+^ and then a Ca^2+^ saturating (5 mmol/L) solutions, both containing 10 μmol/L ionomycin (Sigma, St. Louis, MO) until reaching a minimal (R_*min*_) or a maximal (R_*max*_) ratio values, respectively. [Ca^2+^]_i_ was calculated as described previously using the following equation:

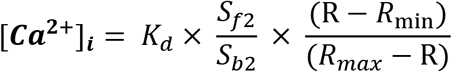

*K_d_* (dissociation constant) in adult myocytes was taken as 224 nmol/L. The scaling factors S_f2_ and S_b2_ were extracted from calibration as previously described^36^. Δ[Ca^+2^]_i_ amplitude was considered as: peak [Ca^+2^]_i_ – resting [Ca^+2^]_i_.

The GHRH-A, MR-409 (N-Me-Tyr1, D-Ala2, Orn12,21, Abu15, Nle27, Asp28)-GHRH(1-29) NHCH_3_) was synthetized by solid phase methods and purified by HPLC, as described previously^5^.

### Statistics

Data distribution was assessed with Pearson normality test. Two-way ANOVA and multiple comparisons were estimated using the Bonferroni and Turkey corrections and expressed as Mean ± SE for normally distributed data. Mann-Whitney test was used to evaluate continuous variables non-normally distributed and described by median and interquartile range [IQR]. All statistics were tested using two-sided at alpha=0.05. (GraphPad Software, Inc. La Jolla, CA).

## RESULTS

The renal embolization was performed in 36 Yorkshire swine. Sixteen animals successfully completed the study. Twenty animals died before initiation of the treatment, the majority in the first 2 initial weeks from which: 1 was euthanized because seizures and 1 because of uremic encephalopathy that did not responded to treatment. Three had complications during the embolization procedure because of arteriovenous malformation, myocardial infarction, and anaphylaxis, respectively. After the procedure, 1 animal died as a complication of a central line placement, 12 died secondary to acute kidney injury. One animal died at 12-weeks during the electrical stimulation procedure, which reproduced a ventricular tachycardia that converted to ventricular fibrillation that did not respond to treatment.

CKD was confirmed 12-weeks post-embolization by increased creatinine (Δ1.09±0.13; CI95% 0.82-1.36; p<0.0001) (Figure 1A) and BUN (Δ10.38±1.313; CI95% 7.58-13.17; p<0.0001) (Figure 1B). Hemoglobin decreased [9.95 (9.25, 11.15) to 7.15 (6.5, 7.85 p<0.0001)] (Figure 1C). and mean arterial pressure (MAP) increased by 7.43±3.31 (CI95% 0.38-14.48; p=0.04) (Figure 1D).

**Figure 1.**
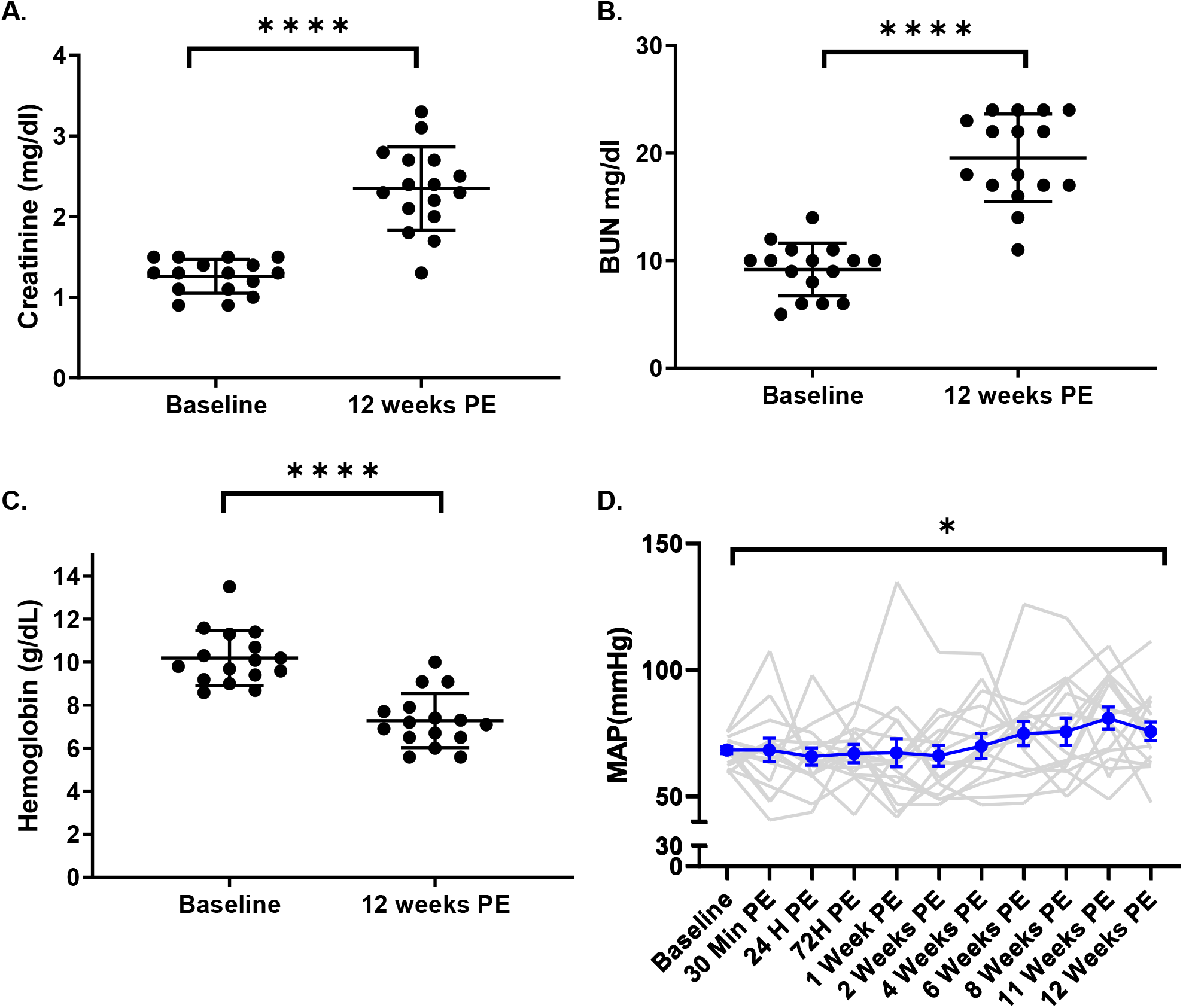
Development of Chronic Kidney Disease at 12 weeks post-embolization. Yorkshire swine (n=16) developed signs of CKD after catheter-guided renal artery embolization producing a 5/6 nephrectomy, evidenced by increased **A**. creatinine (p<0.0001), **B**. Blood urea nitrogen (BUN, p<0.0001), and **C**. mean arterial pressure (MAP, p=0.04), and decreased **D**. hemoglobin (p<0.0001).

HFpEF was evident at 12-weeks by increased left ventricular end-diastolic (LVED)-Mass corrected by body surface area (BSA) [baseline: 81.84 (78.65, 88.83), 12-weeks: 107.0 (93.56, 148.9; p=0.0002; Figure 2], elevated relative wall thickness (RWT) by 0.120±0.019 (CI95% 0.08-0.161; p<0.0001; Figure 2B-C), end-diastolic pressure (EDP) by 8.97±2.40 (CI95% 3.84-14.10; p=0.002; Figure 3A), EDP/end-diastolic volume (EDV) ratio/BSA by 0.09±0.02 (p=0.002; Figure 3B), and tau by 23.53±5.756 (CI95% 11.26 to 35.80; p=0.001; Figure 3C). Ejection fraction (EF) was maintained within the normal range throughout the study. One animal reproduced ventricular tachycardia after electrical stimulation and was not revertible at 12 weeks.

**Figure 2.**
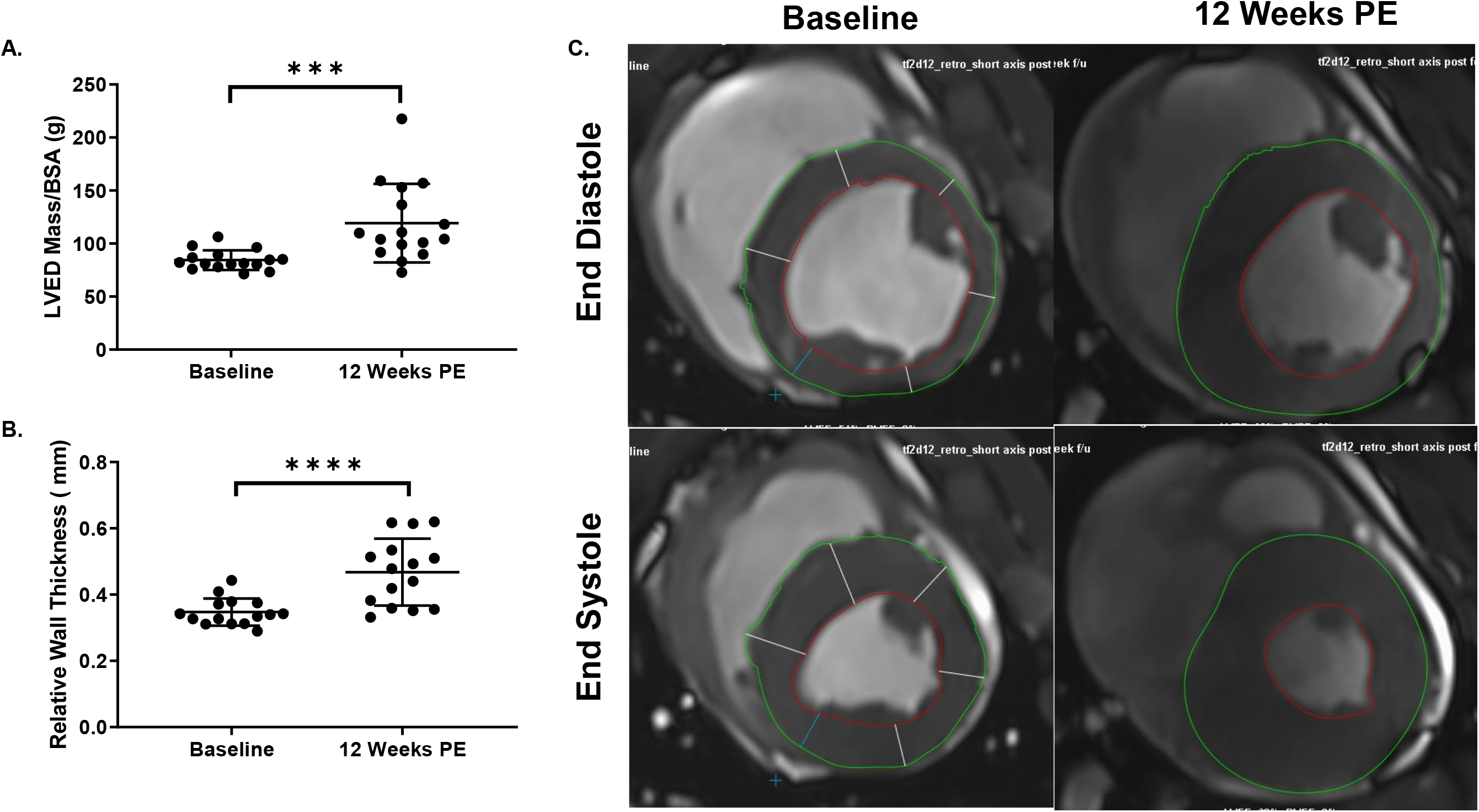
Left ventricular hypertrophy at 12 weeks post-embolization. CKD-induced left ventricular hypertrophy evidenced by **A**. left ventricular end-diastolic (LVED)-Mass corrected by body surface area (BSA) (p=0.0002), **B**. relative wall thickness (RWT; p<0.0001), and **C**. representative MRI at baseline and 12 weeks post-embolization in end diastole and end systole.

**Figure 3.**
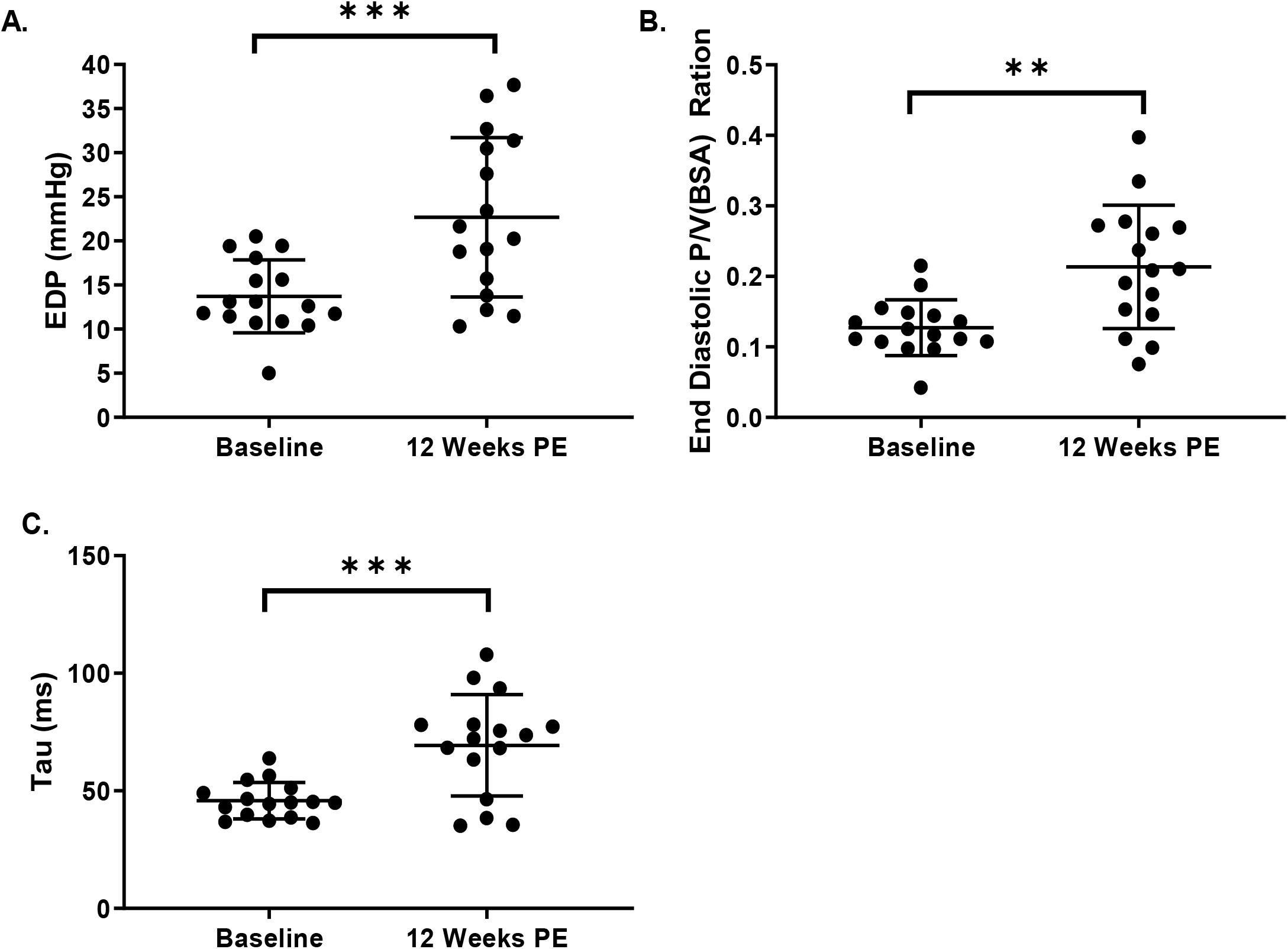
Hemodynamic HFpEF parameters at 12 weeks post-embolization. Features of HFpEF were evidenced by increased **A**. end-diastolic pressure (EDP; p=0.002), **B**. EDP/end-diastolic volume (EDV) ratio/BSA (0.002), and **C**. tau by (p=0.001).

### Response to GHRH

At study completion (16-18 weeks), BUN was significantly lower by 6.62±2.3mg/dL (CI95% 0.89-12.36; p=0.019) in GHRH-A compared to placebo treated animals. Creatinine, MAP, total kidney mass, anemia, and EF remained stable (Table 1).

**TABLE 1.**

| **PARAMETER** | **GROUP** | **BASELINE** | **12 WEEKS POST-**<br>**EMBOLIZATION** | **16-18 WEEKS**<br>**POST-**<br>**EMBOLIZATION** | **2-WAY**<br>**ANOVA**<br>**P VALUE** |
| --- | --- | --- | --- | --- | --- |
| **MAP**<br>**(MMHG)** | **Placebo** | 70.32±1.47 | 77.39±3.54 | 73.54±4.19 | 0.94 |
|  | **GHRH-A** | 65.93±2.07 | 73.72±6.75 | 71.49±7.40 |  |
| **CREATININE**<br>**(MG/DL)** | **Placebo** | 1.313±0.04 | 2.59±0.12 | 2.67±0.22 | 0.27 |
|  | **GHRH-A** | 1.213±0.09 | 2.11±0.2 | 2.17±0.21 |  |
| **BUN (MG/DL)** | **Placebo** | 9.25±0.61 | 21.75±0.98 | 22.88±2.81 | 0.11 |
|  | **GHRH-A** | 9.125±1.11 | 17.38±1.46 | 16.25±1.83 |  |
| **HGB**<br>**(MG/DL)** | **Placebo** | 10.25±0.23 | 7.26±0.58 | 8.35±0.3 | 0.81 |
|  | **GHRH-A** | 10.14±0.62 | 7.31±0.29 | 7.66±0.34 |  |
| **HTC (%)** | **Placebo** | 33 (32,36) | 22.5 (19,29.25) | 27 (24,30.25) | 0.89 |
|  | **GHRH-A** | 30 (28.25,38) | 21.5 (21,24.5) | 23.5 (22.25,23.5) |  |

In the GHRH-A group, diastolic evaluation revealed reduced EDP by −9.34±3.82mmHg (CI95% 0.88-17.80; p= 0.027), associated with recovered passive diastolic function and decreased (EDP/EDV)/BSA by 0.094 (CI95% 0.01-0.17; p=0.018; Figure. 4A-B). Active relaxation, characterized by tau (Figure. 4C-D), and isovolumetric relaxation, measured by dP/dtmin, were both improved in the GHRH-A group, but were not significantly different between groups. Stroke work (SW) increased over time in the placebo group by 1482 mmHg*mL (CI95% 2370-593.8; p=0.0007; Figure. 4E). Preload recruitable stroke work (PRSW) and dP/dtmax were similar between groups with no changes over time. The percent decrease in LVED-Mass/BSA was greater (albeit not significantly) in the GHRH-A group compared to placebo (−11.79±6.423%; −25.57-1.984; p=0.08; Fig. 4F) (Table 2). RWT (Figure.4G), EDV, ESV, stroke volume (SV) corrected by BSA, and dry lung weight were similar between the two groups. Importantly, ventricular tachycardia was reproduced by electrical stimulation in one animal of the placebo group and none of the GHRH group. The calcium transient amplitude was increased (p=0.009) in cardiomyocytes from GHRH-A-treated animals, although the rate of calcium decay was not affected (Figure. 4H).

**Figure 4.**
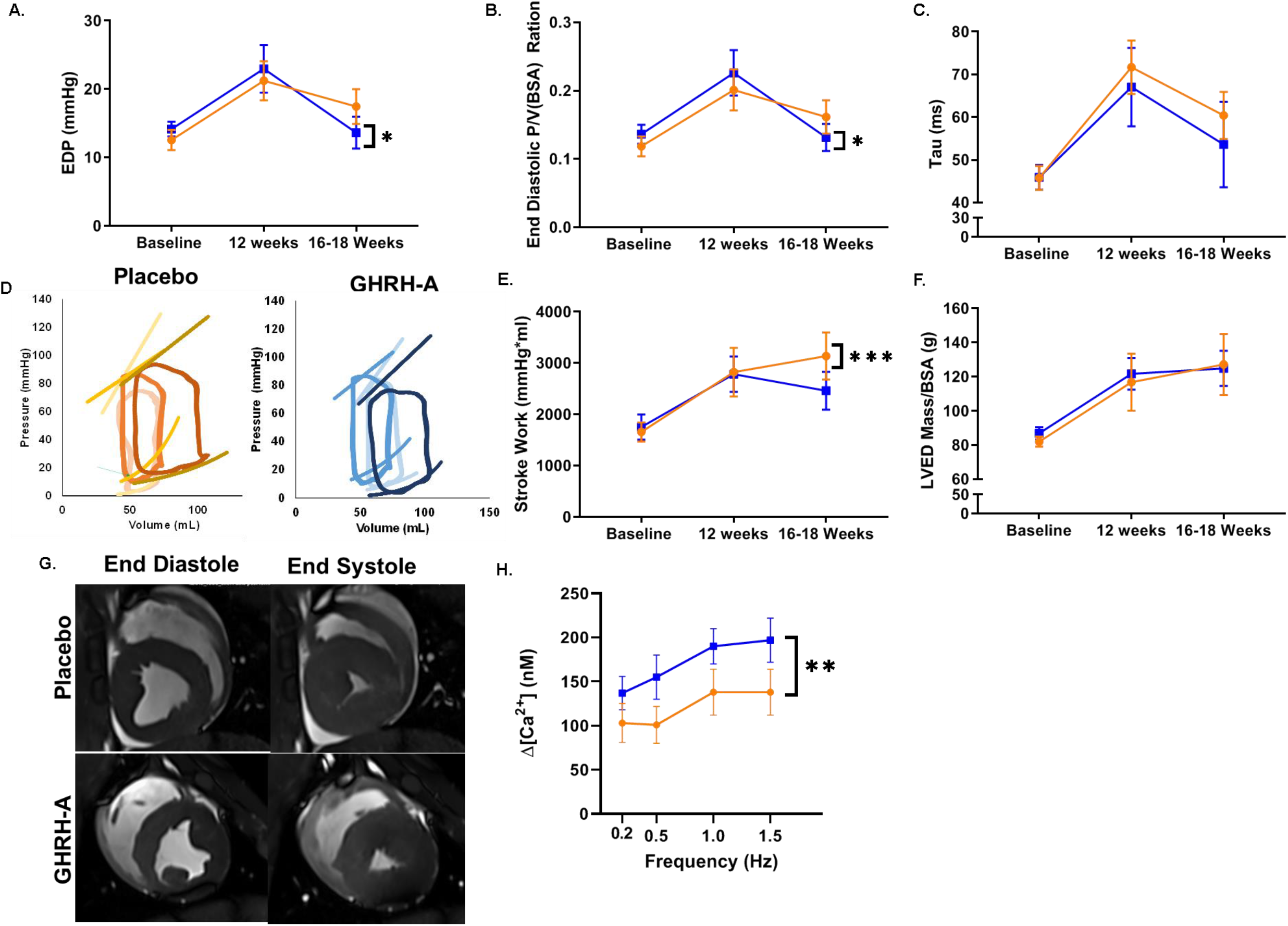
Effectiveness of GHRH-A in a large animal model of HFpEF. GHRH-A improves diastolic function including: **A)** end diastolic pressure (EDP), **B)** EDP/end-diastolic volume corrected by BSA and **C)** Tau. **D)** Representative PV loops (GHRH-A: Blue, Placebo: Orange; Baseline: light color, 12 Weeks PE: bright color; 18 Weeks PE: dark color). CKD-induced HFpEF phenotype presents with increased **E)** stroke work, **F)** left ventricular end-diastolic (LVED) mass/BSA, and **G)** relative wall thickness (RWT). **(H)** Cardiomyocyte calcium-transient amplitude (p=0.009).

**TABLE 2.**

| PARAMETER | GROUP | BASELINE | 12 WEEKS POST-EMBOLIZATION | 16-18 WEEKS POST-EMBOLIZATION | 2-WAY ANOVA<br>P VALUE |
| --- | --- | --- | --- | --- | --- |
| *EF (%)* | Placebo | 41.95±2.46 | 49.65±2.64 | 50.63±3.02 | 0.52 |
|  | GHRH-A | 40.23±1.91 | 50.63±2.25 | 51.64±3.42 |  |
| *EDV/BSA (ML)* | Placebo | 107±3.03 | 109.7±6.95 | 110.1±5.26 | 0.94 |
|  | GHRH-A | 105.9±4.38 | 101.5±4.68 | 101.7±4.09 |  |
| *ESV/BSA (ML)* | Placebo | 62.08±3.15 | 54.62±3.16 | 54.69±5.00 | 0.23 |
|  | GHRH-A | 63.44±3.72 | 50.4±3.85 | 49.31±4.36 |  |
| *SV/BSA (ML)* | Placebo | 43.51 (38.15, 50.55) | 50.51 (45.54, 57.45) | 53.82 (47.81, 57.58) | 0.08 |
|  | GHRH-A | 41.93 (37.12, 49.56) | 51.81 (44.75, 56.34) | 49.31 (43.91, 59.12) |  |
| *LVED*<br>*MASS/BSA (G)* | Placebo | 82.01±2.93 | 116.8±16.72 | 127.2±17.9 | 0.86 |
|  | GHRH-A | 86.88±3.59 | 121.7±9.31 | 124.9±10.33 |  |
| *RELATIVE*<br>*WALL*<br>*THICKNESS* | Placebo | 0.330±0.01 | 0.44±0.03 | 0.49±0.05 | 0.86 |
|  | GHRH-A | 0.362±0.02 | 0.49±0.04 | 0.54±0.04 |  |
| *EDP MMHG* | Placebo | 12.47±1.52 | 21.12±2.87 | 17.44±2.58 | 0.42 |
|  | GHRH-A | 14.21±1.05 | 23.01±3.47 | 13.63±2.34 |  |
| *P/V*<br>*RATIO/BSA* | Placebo | 0.119±0.01 | 0.2±0.03 | 0.16±0.02 | 0.43 |
|  | GHRH-A | 0.136±0.01 | 0.23±0.03 | 0.13±0.02 |  |
| *EDPVR* | Placebo | 0.227±0.04 | 0.26±0.04 | 0.26±0.03 | 0.37 |
|  | GHRH-A | 0.252±0.02 | 0.41±0.10 | 0.22±0.03 |  |
| *TAU* | Placebo | 45.47(37.59, 53.85) | 74.63 (65.5, 77.96) | 58.54 (49.1, 76.51) | 0.88 |
|  | GHRH-A | 44.97(39.8, 48.09) | 68.21 (40.44, 89.66) | 49.16 (37.44, 58.98) |  |
| *ESPVR* | Placebo | 0.936±0.16 | 0.6±0.06 | 0.65±0.07 | 0.16 |
|  | GHRH-A | 0.964±0.11 | 0.9±0.17 | 0.5±0.08 |  |
| *PRSW* | Placebo | 0.944±0.35 | 5.28±1.93 | 2.0±1.01 | 0.96 |
|  | GHRH-A | 0.957±0.28 | 4.76±2.32 | 1.5±0.75 |  |
| *DP/DTMAX* | Placebo | 841.2 (672.3, 918.4) | 813.4 (693.3, 945.5) | 867.1 (625.3, 1052) | 0.32 |
|  | GHRH-A | 764 (678.5, 916.8) | 990.2 (746.7, 1248) | 854.3 (472.2, 955.1) |  |

## DISCUSSION

This blinded, placebo-controlled, preclinical study tested the effectiveness at improving the HFpEF phenotype of a synthetic, potent GHRH-A compared to placebo in a large animal model of CKD-induced HFpEF. The encouraging results demonstrate that GHRH-A produced a substantial improvement in cardiac diastolic function in animals with established HFpEF. Hypertrophy of the left ventricle and SW elevation were evident, similar to that described in the CKD-HFpEF phenotype^4^. Ventricular tachycardia was stimulated in two animals, corroborating structural (pro-arrhythmic) changes in the myocardium. Together these preclinical findings support ongoing translational development of this class of agents for human HFpEF.

GHRH-A reparative effects on HFpEF likely occur through multiple mechanisms^15^, including anti-apoptotic effects via inhibition of ERK1/2 and PI3K/Akt signaling associated with elevated Blc-2 and reduced Bax:Bcl-2^6^, stimulation of cardiac progenitor proliferation^6, 16^, inhibition of hypertrophy, including Gq signaling and downstream pathways^17^, reduced inflammation, a major HFpEF-characteristic, specifically plasma levels of IL-2, IL-6 and TNF-α^6^, reduced expression of pro-fibrotic pathways genes^6^, and better organized collagen deposition^6^. In a mouse model of HFpEF, GHRH-A inhibited cardiomyocyte sarcomere and contractile dysfunction, preventing Ca^2+^ leak from the sarcoplasmic reticulum and improving sarcomere relaxation and calcium handling^9^. The anti-arrhythmogenic effect, independent of the scar reduction seen in GH therapy^18^, was also seen with this GHRH-analogue, a poorly understood effect. To our knowledge, this is the first study evaluating GHRH-A in a large animal model of HFpEF. GHRH-A improved diastolic dysfunction, supporting the feasibility and safety of this approach and suggesting its potential therapeutic application for HFpEF.

## REFERENCES

1. Pfeffer MA, Shah AM and Borlaug BA. Heart Failure With Preserved Ejection Fraction In Perspective. Circulation Research. 2019;124:1598–1617.

2. Pieske B, Tschöpe C, de Boer RA, Fraser AG, Anker SD, Donal E, Edelmann F, Fu M, Guazzi M, Lam CSP, Lancellotti P, Melenovsky V, Morris DA, Nagel E, Pieske-Kraigher E, Ponikowski P, Solomon SD, Vasan RS, Rutten FH, Voors AA, Ruschitzka F, Paulus WJ, Seferovic P and Filippatos G. How to diagnose heart failure with preserved ejection fraction: the HFA–PEFF diagnostic algorithm: a consensus recommendation from the Heart Failure Association (HFA) of the European Society of Cardiology (ESC). European Heart Journal. 2019;40:3297–3317.

3. Lam CSP, Voors AA, de Boer RA, Solomon SD and van Veldhuisen DJ. Heart failure with preserved ejection fraction: from mechanisms to therapies. European Heart Journal. 2018;39:2780–2792.

4. Shah SJ, Katz DH, Selvaraj S, Burke MA, Yancy CW, Gheorghiade M, Bonow RO, Huang CC and Deo RC. Phenomapping for novel classification of heart failure with preserved ejection fraction. Circulation. 2015;131:269–79.

5. Cai R, Schally AV, Cui T, Szalontay L, Halmos G, Sha W, Kovacs M, Jaszberenyi M, He J, Rick FG, Popovics P, Kanashiro-Takeuchi R, Hare JM, Block NL and Zarandi M. Synthesis of new potent agonistic analogs of growth hormone-releasing hormone (GHRH) and evaluation of their endocrine and cardiac activities. Peptides. 2014;52:104–12.

6. Kanashiro-Takeuchi RM, Szalontay L, Schally AV, Takeuchi LM, Popovics P, Jaszberenyi M, Vidaurre I, Zarandi M, Cai R-Z, Block NL, Hare JM and Rick FG. New therapeutic approach to heart failure due to myocardial infarction based on targeting growth hormone-releasing hormone receptor. Oncotarget. 2015;6:9728–9739.

7. Kanashiro-Takeuchi RM, Takeuchi LM, Rick FG, Dulce R, Treuer AV, Florea V, Rodrigues CO, Paulino EC, Hatzistergos KE, Selem SM, Gonzalez DR, Block NL, Schally AV and Hare JM. Activation of growth hormone releasing hormone (GHRH) receptor stimulates cardiac reverse remodeling after myocardial infarction (MI). Proceedings of the National Academy of Sciences of the United States of America. 2012;109:559–63.

8. Bagno LL, Kanashiro-Takeuchi RM, Suncion VY, Golpanian S, Karantalis V, Wolf A, Wang B, Premer C, Balkan W, Rodriguez J, Valdes D, Rosado M, Block NL, Goldstein P, Morales A, Cai R-Z, Sha W, Schally AV and Hare JM. Growth hormone-releasing hormone agonists reduce myocardial infarct scar in swine with subacute ischemic cardiomyopathy. Journal of the American Heart Association. 2015;4:e001464.

9. Dulce RA, Kanashiro-Takeuchi RM, Takeuchi LM, Salerno AG, Kulandavelu S, Balkan W, Zuttion MSSR, Cai R, Schally AV and Hare JM. Synthetic Agonist of Growth Hormone-Releasing Hormone as Novel Treatment for Heart Failure with Preserved Ejection Fraction. bioRxiv. 2020:2020.02.28.967000.

10. Siskin GP, Dowling K, Virmani R, Jones R and Todd D. Pathologic Evaluation of a Spherical Polyvinyl Alcohol Embolic Agent in a Porcine Renal Model. Journal of Vascular and Interventional Radiology. 2003;14:89–98.

11. Misra S, Gordon JD, Fu AA, Glockner JF, Chade AR, Mandrekar J, Lerman L and Mukhopadhyay D. The porcine remnant kidney model of chronic renal insufficiency. The Journal of surgical research. 2006;135:370–9.

12. Williams AR, Hatzistergos KE, Addicott B, McCall F, Carvalho D, Suncion V, Morales AR, Da Silva J, Sussman MA, Heldman AW and Hare JM. Enhanced effect of combining human cardiac stem cells and bone marrow mesenchymal stem cells to reduce infarct size and to restore cardiac function after myocardial infarction. Circulation. 2013;127:213–23.

13. Natsumeda M, Florea V, Rieger AC, Tompkins BA, Banerjee MN, Golpanian S, Fritsch J, Landin AM, Kashikar ND, Karantalis V, Loescher VY, Hatzistergos KE, Bagno L, Sanina C, Mushtaq M, Rodriguez J, Rosado M, Wolf A, Collon K, Vincent L, Kanelidis AJ, Schulman IH, Mitrani R, Heldman AW, Balkan W and Hare JM. A Combination of Allogeneic Stem Cells Promotes Cardiac Regeneration. J Am Coll Cardiol. 2017;70:2504–2515.

14. Coppini R, Ferrantini C, Aiazzi A, Mazzoni L, Sartiani L, Mugelli A, Poggesi C and Cerbai E. Isolation and functional characterization of human ventricular cardiomyocytes from fresh surgical samples. Journal of visualized experiments: JoVE. 2014.

15. Schally AV, Zhang X, Cai R, Hare JM, Granata R and Bartoli M. Actions and Potential Therapeutic Applications of Growth Hormone-Releasing Hormone Agonists. Endocrinology. 2019;160:1600–1612.

16. Florea V, Majid SS, Kanashiro-Takeuchi RM, Cai RZ, Block NL, Schally AV, Hare JM and Rodrigues CO. Agonists of growth hormone-releasing hormone stimulate self-renewal of cardiac stem cells and promote their survival. Proceedings of the National Academy of Sciences of the United States of America. 2014;111:17260–5.

17. Gesmundo I, Miragoli M, Carullo P, Trovato L, Larcher V, Di Pasquale E, Brancaccio M, Mazzola M, Villanova T, Sorge M, Taliano M, Gallo MP, Alloatti G, Penna C, Hare JM, Ghigo E, Schally AV, Condorelli G and Granata R. Growth hormone-releasing hormone attenuates cardiac hypertrophy and improves heart function in pressure overload-induced heart failure. Proceedings of the National Academy of Sciences of the United States of America. 2017;114:12033–12038.

18. Stamatis KV, Kontonika M, Daskalopoulos EP and Kolettis TM. Electrophysiologic Effects of Growth Hormone Post-Myocardial Infarction. International journal of molecular sciences. 2020;21.

